# Mitochondrial variant enrichment from high-throughput single-cell RNA-seq resolves clonal populations

**DOI:** 10.1101/2021.03.08.434450

**Authors:** Tyler E. Miller, Caleb A. Lareau, Julia A. Verga, Daniel Ssozi, Leif S. Ludwig, Chadi El Farran, Gabriel K. Griffin, Andrew A. Lane, Bradley E. Bernstein, Vijay G. Sankaran, Peter van Galen

**Affiliations:** Department of Pathology, Massachusetts General Hospital, Boston, MA; Broad Institute of MIT and Harvard, Cambridge, MA; Department of Pathology, Stanford University, Palo Alto, CA; Division of Hematology, Brigham and Women’s Hospital, Boston, MA; Division of Hematology/Oncology, Boston Children’s Hospital and Department of Pediatric Oncology, Dana-Farber Cancer Institute, Harvard Medical School, Boston, MA; Berlin Institute of Health at Charité - Universitätsmedizin Berlin, Berlin Institute for Medical Systems Biology, Max Delbrück Center for Molecular Medicine in the Helmholtz Association, Berlin, Germany; Department of Pathology, Brigham and Women’s Hospital, Boston, MA; Department of Medical Oncology, Dana-Farber Cancer Institute, Boston, MA; Ludwig Center at Harvard, Harvard Medical School, Boston, MA

## Abstract

Reconstructing lineage relationships in complex tissues can reveal mechanisms underlying development and disease. Recent methods combine single-cell transcriptomics with mitochondrial DNA variant detection to establish lineage relationships in primary human cells, but are not scalable to interrogate complex tissues. To overcome this limitation, here we develop a technology for high-confidence detection of mitochondrial mutations from high-throughput single-cell RNA-sequencing. We use the new method to identify skewed immune cell expansions in primary human clonal hematopoiesis.

## Main text

Single-cell RNA-sequencing (scRNA-seq) enables the unbiased assessment of cell states in health and disease^1,2^. Combined acquisition of cell state and genetic information can provide additional insight, such as targeted enrichment of cancer driver mutations from single-cell transcriptomes^3,4^. Separately, combining scRNA-seq with genetic cell barcodes is a powerful method to reveal clonal relationships and evolutionary dynamics of cells within organisms^5,6^. However, this has largely been limited to experimental model systems that can be genetically manipulated to insert cell barcodes. To infer clonal dynamics in primary human cells, recent methods have detected and utilized mitochondrial DNA (mtDNA) mutations as naturally occurring genetic cell barcodes^7–9^. The combination of scRNA-seq with mtDNA mutation detection can inform clonal relationships with high confidence, but is currently restricted to expensive, low-throughput, full-length transcript sequencing technologies like SmartSeq2^7,10^. To enable the reconstruction of clonal relationships in complex human tissues, we developed a method that captures genetic variants from high-throughput scRNA-seq platforms: MAESTER, or Mitochondrial Alteration Enrichment from Single-cell Transcriptomes to Establish Relatedness (Figure 1A). MAESTER is compatible with the most common high-throughput scRNA-seq platforms, including 10x Genomics 3’ protocols, Seq-Well S^3^, and Drop-seq (Supplemental Figures 1-3)^11,12^. An intermediate step in each of these platforms yields full length cDNA transcripts, from which we enrich all 15 mitochondrial transcripts using pools of primers, while maintaining cell-identifying barcodes (Figure 1B, Supplemental Figure 4). Standard next-generation sequencing with 250 bp reads is then used to obtain the sequence of the amplified mitochondrial transcripts (Figure 1A). We developed a computational toolkit to call mtDNA variants from MAESTER data, the Mitochondrial Alteration Enrichment and Genome Analysis Toolkit (maegatk, Supplemental Figure 5, Methods). Building on previous tools that we developed^8^ for mtDNA variant detection from single-cell ATAC or SmartSeq2, maegatk specifically handles technical biases implicit in high-throughput transcriptomic libraries. Critically, maegatk leverages unique molecular identifiers (UMIs) to collapse multiple sequencing reads of the same starting transcript, creating a consensus call for every nucleotide based on the most common call and base quality. This approach mitigates sequence errors introduced during PCR and sequencing and is essential to obtain high-confidence variant calls from high-throughput scRNA-seq protocols. Alterations in mtDNA are then used to infer relatedness between cells.

**Figure 1.**
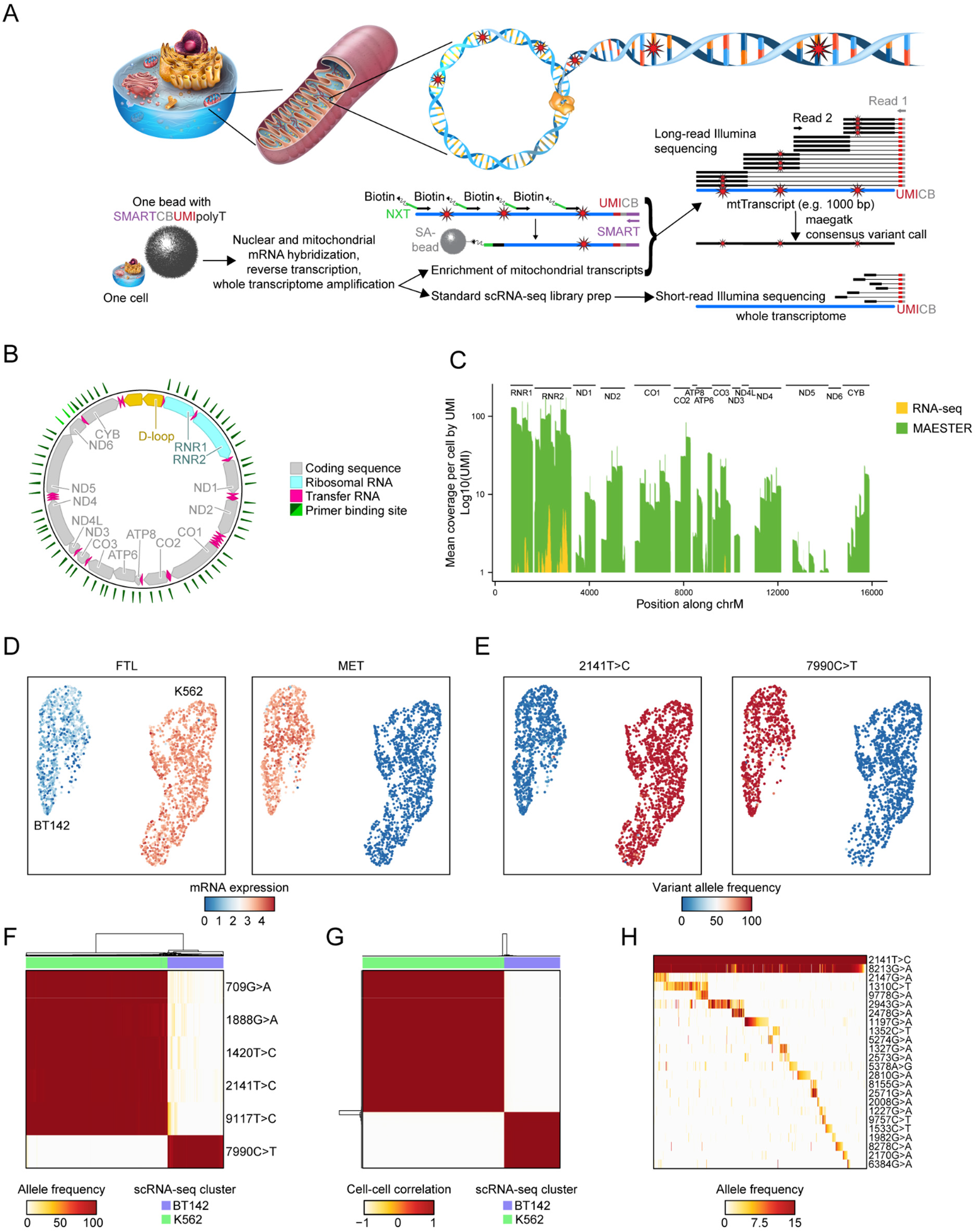
Targeted enrichment of mitochondrial transcripts enables discrimination between genetic clones. **A**. Schematic shows the procedures for lineage inference from single-cell transcriptomes using MAESTER. **B**. Diagram depicts the circular mitochondrial genome with annotated features. The green triangles indicate where MAESTER primers bind. **C**. Bar plot shows coverage of the mitochondrial genome with and without amplification from Seq-Well libraries using MAESTER. Mean coverage of 2,482 K562 and BT142 cells is shown. Each of the transcripts (UMIs) was sequenced ≥3 times. **D-E**. UMAPs show detection of cell type-specific (D) gene expression from scRNA-seq and (E) homoplasmic mtDNA variants from MAESTER. Only cells with >3-fold coverage of the indicated variants are shown. **F**. Heatmap depicts separation of 1,523 K562 and BT142 cells (columns) based on six mtDNA variants (rows). Cell type annotation from scRNA-seq is shown on top. **G**. Correlation matrix shows cell similarity based on the allele frequencies of six homoplasmic variants (rows and columns depict 1,523 cells). Unsupervised clustering identified two clusters that correlate with cell annotations from scRNA-seq. For F and G, only cells with >3-fold coverage of each homoplasmic variant are shown. **H**. Heatmap shows VAF of 23 mtDNA variants detected by MAESTER (rows) for 596 K562 cells (columns) with informative subclonal variants. Homoplasmic K562 variant 2141T>C is shown for comparison. Heatmap is organized by clonal structure (Methods).

We established the feasibility and efficiency of this method using human cell mixing experiments. Chronic myelogenous leukemia cells (K562) were mixed with brain tumor cells (BT142) and analyzed using standard high-throughput scRNA-seq runs with Seq-Well S^3^ and 10x 3’ v3. MAESTER dramatically increased the coverage of mitochondrial transcripts compared to scRNA-seq data alone (mean coverage per cell 0.2 to 0.7-fold for RNA-seq, 52 to 217-fold for MAESTER, Figure 1C, Supplemental Figure 6A-B), enabling reliable mtDNA variant calling with high confidence across many of the transcripts. Using Uniform Manifold Approximation and Projection (UMAP) for dimensionality reduction, the two cell populations cluster based on mRNA expression data (Figure 1D). MAESTER enabled identification of six homoplasmic mtDNA variants that distinguished between cell types (Figure 1E-G, Supplemental Figure 6C-D). Combining data from all six informative variants cleanly separates cell types and demonstrated 100% concordance with mRNA clusters (Supplemental Figure 6E-F). Of note, MAESTER identified the same six variants in the Seq-Well and 10x libraries (Supplemental Figure 7A-D).

To benchmark MAESTER’s ability to identify clonal structure at a more granular resolution, we performed a clonal expansion experiment. One hundred K562 cells were plated and allowed to expand for 14 days (doubling time ∼24h), followed by scRNA-seq with MAESTER. We identified 23 informative mtDNA variants that revealed clonally related populations of K562 cells (Figure 1H, Supplemental Figure 7E), demonstrating the capacity of this method to resolve subpopulations within closely related cells.

We next applied MAESTER to derive clonal structure within primary human patient specimens. We utilized a bone marrow aspirate from a patient with *TET2*-mutated clonal hematopoiesis. The clonal hematopoiesis had evolved into blastic plasmacytoid dendritic cell neoplasm (BPDCN), as the patient had skin tumors at the time of collection. However, the concurrent bone marrow aspirate we utilized showed no tumor involvement (Methods). We performed 10x single-cell sequencing with MAESTER on this bone marrow aspirate and identified 9,346 high-quality cells, including all expected cell types, with an abundance of cytotoxic T-cells (CTLs), possible related to his evolving malignancy (CTLs, Figure 2A, Supplemental Figure 8A). We identified 23 clones readily distinguishable from each other using 26 informative mtDNA variants (1,397 cells were assigned to a clone, Figure 2B), indicating MAESTER can resolve clonal populations in primary human specimens.

**Figure 2.**
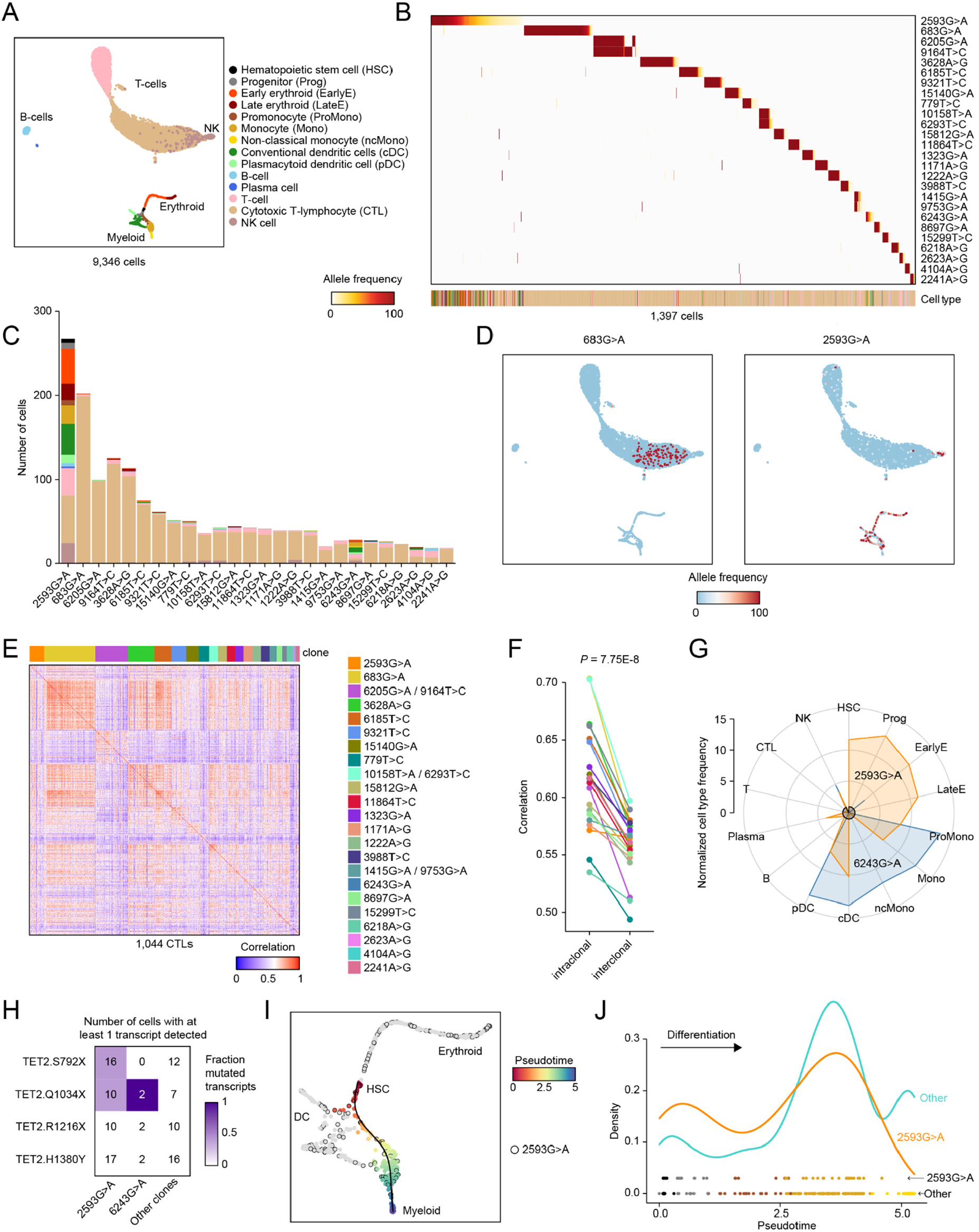
Genetic clones exhibit lineage bias in clonal hematopoiesis. **A.** UMAP of all 9,346 cells profiled by 10x scRNA-seq from a bone marrow aspirate from an individual with clonal hematopoiesis. **B**. Heatmap shows VAF of 26 informative mtDNA variants detected by MAESTER and maegatk (rows) for 1,397 cells (columns) with at least 1% VAF of one of the 26 variants. Heatmap is organized by clones and sorted by clone size. Cell type is listed on the bottom by color according to legend in A. **C**. Stacked bar graph of the number of cells in each clone, with cell type denoted by color according to legend in A. **D**. UMAPs display VAF in each cell for mtDNA variants 683G>A (left) and 2593G>A (right). **E**. Heatmap shows gene expression correlation of 1,044 CTLs in mitochondrial-variant defined clones (all CTLs from B and C). Color indicates Pearson correlation between expression of 100 variable genes across CTLs. **F**. Paired line graph plotting average cell-cell correlation for all CTLs within each clone (intraclonal) and average cell-cell correlation for all CTLs from one clone to all other CTLs. *P*-value calculated by paired Student’s t-Test. Clones are colored as in E. **G**. Radar plot shows normalized cell type frequencies for clones marked by 2593G>A or 6243G>A. Frequency is normalized to the frequency of cell types for all cells, which is set to 1 and denoted by black circle at the center. **H**. Chart with number of cells that have at least one wild-type or mutated transcript detected by GoT for the four TET2 mutations known to be present in the patient’s cells. Fraction of mutated transcripts for all cells in each box designated by color scale. **I**. UMAP depicts pseudotime of myeloid differentiation. Cells in the 2593G>A clone are marked by a black circle. **J**. Line graph shows relative cell density along pseudotime of myeloid differentiation in clone 2593G>A vs. all other myeloid cells. Symbols at the bottom indicate the cells, colored according to legend in A.

We found that 9/23 clones were skewed towards CTLs (Figure 2C, Supplemental Figure 8B-C). Interestingly, most of the CTL-biased clones clustered together in the UMAP (Figure 2D, Supplemental Figure 9). Indeed, we found that cells within the clones were transcriptionally more similar to each other than to other CTL-biased clones (*P* = 7.75E-8, Figure 2E, F). This indicates that MAESTER accurately classified these cells as related and providing potential insight into the biology of clonal CTL expansions in the context of a malignancy.

We also identified two clones with significant myeloid lineage bias, identified by mtDNA alterations 2593G>A and 6243G>A (Figure 2C, D, G, Supplemental Figure 8, 9). Given the patient’s clonal hematopoiesis, we sought to understand if these represented expanded *TET2* clones. We utilized Genotyping of Transcriptomes (GoT)^3^ to identify the patient’s four known *TET2* loss-of-function mutations in single cells. We found that cells within the two myeloid-biased clones contained a high fraction of mutated transcripts (Figure 2H). Of cells in the 2593G>A clone, 41% and 40% of the *TET2* transcripts detected had TET2.S792X and TET2.Q1034X mutations, respectively. This is consistent with bi-allelic *TET2* inactivation, a recurrent feature in myeloid malignancies^13^. No mutated transcripts were identified in cells from the other 21 clones we identified, providing evidence that the myeloid-biased clones identified by MAESTER represent the patient’s *TET2*-mutated clonal hematopoiesis.

Within the 23 clones (1,397 cells), GoT for *TET2* mutations only captured genotype information, wildtype or mutant, for 123 transcripts in 70 cells (5.0% of cells had at least one *TET2* variant genotyped, most by a single transcript). This relative lack of genotyping efficiency is related to gene expression and amplicon size, and is similar to other protocols that genotype somatic mutations from high-throughput scRNA-seq libraries^4^. In contrast, MAESTER captured the mtDNA genotype at the 2593 position in 21,767 transcripts in 1,396 cells (99.9% of cells, an average of 16 transcripts per cell). However, MAESTER does not capture driver somatic mutations. Thus, when combined, the biological significance of *TET2* mutations detected by GoT and the robust sensitivity of MAESTER allowed us to explore cells marked by 2593G>A as clonally expanded cells with loss of *TET2*.

To interrogate this population further, we compared the myeloid differentiation trajectory of 2593G>A cells to other cells in the bone marrow using pseudotime analysis (Figure 2I)^14^. We found the clonal population was skewed towards less mature cell types (Figure 2J), suggesting an expansion of hematopoietic stem and progenitor cells, consistent with phenotypes observed in *Tet2* knockout mice^15^. The identification and investigation of this clinically relevant pre-malignant population would not have been possible without MAESTER.

In conclusion, MAESTER enables robust mtDNA variant detection in high-throughput 3’-biased scRNA-seq data, which was previously limited to ATAC-seq or full-length scRNA-seq. Due to the widespread use of 10x 3’ protocols, Drop-seq, and Seq-Well, the development of MAESTER makes mtDNA variant detection accessible to more research laboratories and a wider range of experimental contexts. The accompanying maegatk software leverages UMIs to increase confidence in mtDNA variant calls, an advance over previous methods. MAESTER can be implemented on new or prior scRNA-seq datasets by using the amplified cDNA that is stored as a standard practice. The high-throughput nature of 3’-biased scRNA-seq and MAESTER enables the study of clonal relationships and evolutionary dynamics of cells within complex primary human tissues. By developing MAESTER, we democratize and expand the utilization of naturally occurring barcodes created by mitochondrial DNA alterations to discover novel and exciting *in vivo* human biology.

## Acknowledgements

We thank patients for donating bone marrow cells, the Alex Shalek lab, Antonia Kreso and Volker Hovestadt for helpful discussions, and Patricia Rogers and the Broad Institute Flow Facility for technical support. P.v.G., A.A.L., and B.E.B. are supported by the Ludwig Center at Harvard. P.v.G. and V.G.S. are supported by the Harvard Medical School Epigenetics & Gene Dynamics Initiative. P.v.G. is supported by the National Cancer Institute (NCI) R00 Award, Gilead Sciences and the Bertarelli Rare Cancers Fund, and is a Glenn Foundation for Medical Research and AFAR Grant for Junior Faculty awardee. T.E.M. is supported by the American Brain Tumor Association Basic Research Fellowship in honor of Joel A. Gingras, Jr. T.E.M. and J.A.V. are supported by the UK Brain Tumour Charities Future Leaders Award. V.G.S. is supported by the New York Stem Cell Foundation (NYSCF), a gift from the Lodish family to Boston Children’s Hospital, and National Institutes of Health grant R01 DK103794. V.G.S. is a NYSCF-Robertson Investigator.

## Author contributions

T.E.M., C.A.L., J.A.V., D.S. and P.v.G. conducted experiments and analyzed the data. T.E.M., C.A.L., L.S.L., C.E.F., G.K.G., A.A.L., B.E.B., V.G.S. and P.v.G. designed the study and interpreted the data. T.E.M. and P.v.G. wrote the manuscript. All authors edited the manuscript.

## Supplemental data

The supplemental data includes nine figures and two tables. Please contact the corresponding author for the supplemental tables with oligo sequences.

## Supplemental Figures

**Supplemental Figure 1.**
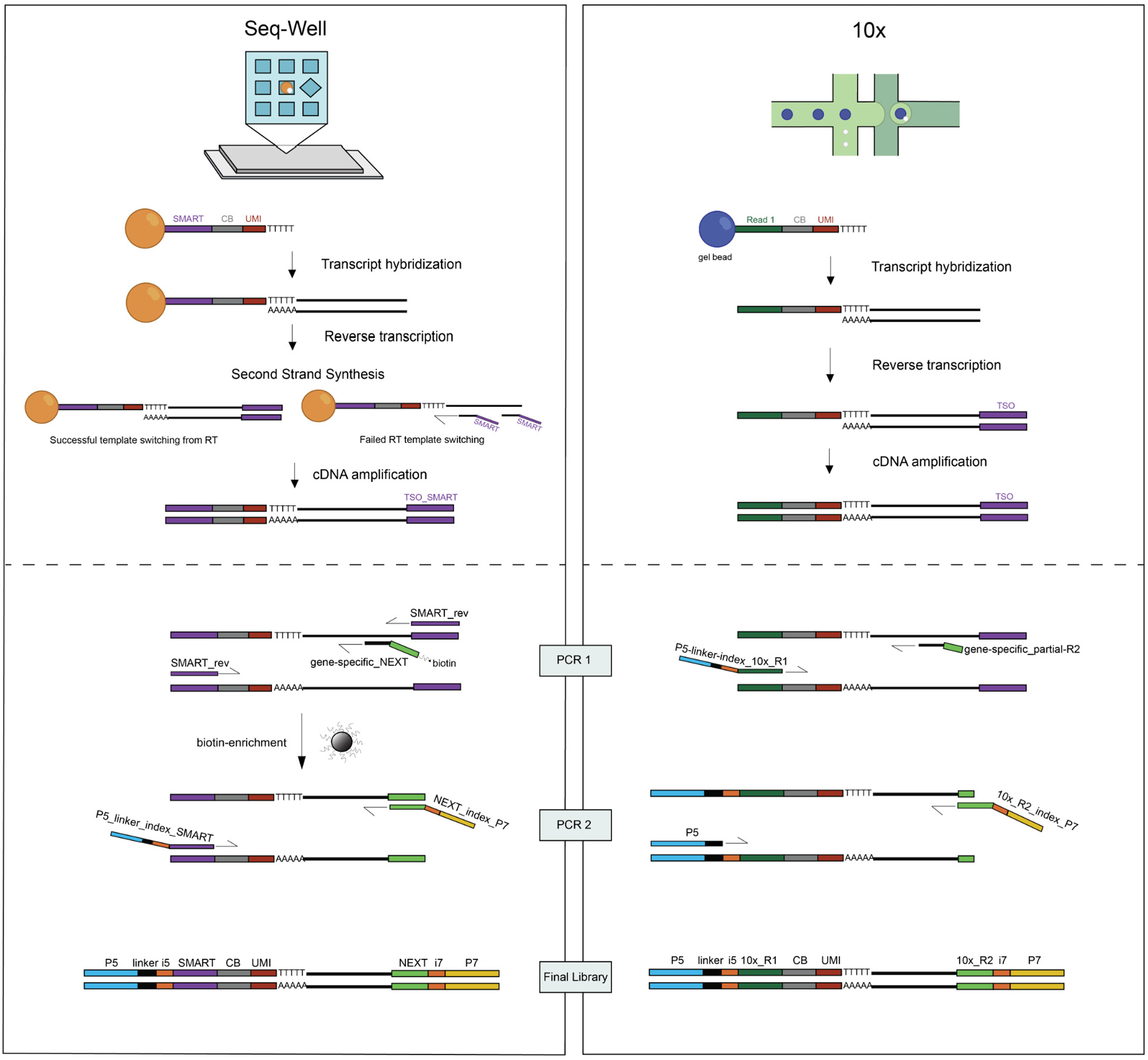
Overview of MAESTER methods for Seq-Well or 10x scRNA-seq. Overview and comparison between the MAESTER method for Seq-Well S^3^ or Drop-Seq scRNA-seq (left) and 10x procedures (right). Above the dotted line is the standard protocol to create the whole transcriptome amplified material (WTA) or cDNA library. Below the dotted line is the mitochondrial enrichment method from the WTA. Note: Seq-Well or Drop-Seq methods use SMART primer sequences on both ends of the cDNA transcripts, necessitating a biotin enrichment step. 10x uses different primers at each end, allowing for an asymmetric PCR without the need for biotin enrichment. CB: cell barcode, UMI: unique molecular identifier.

**Supplemental Figure 2.**
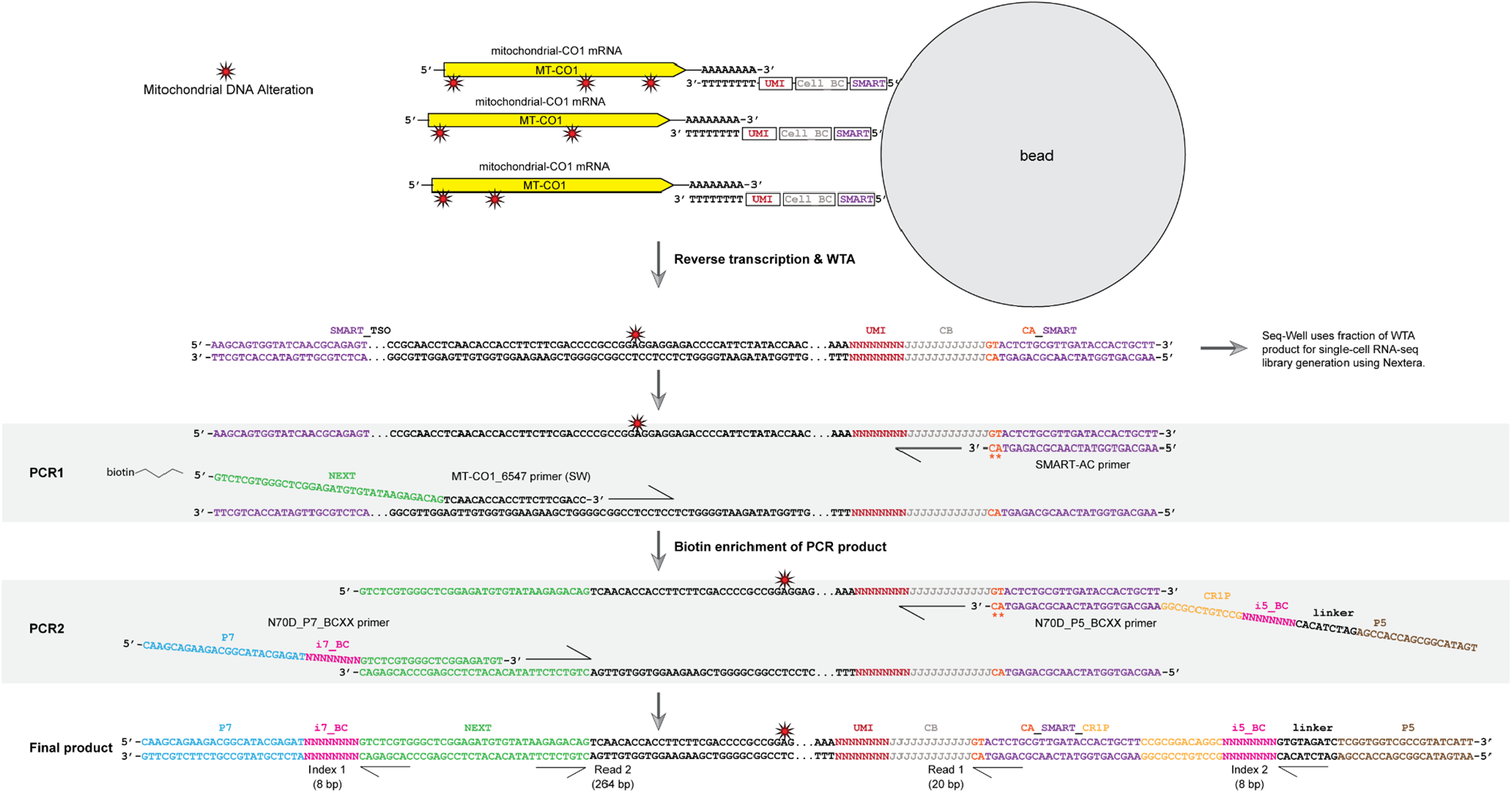
Detailed schematic for MAESTER dial-out PCR protocol from Seq-Well or Drop-Seq WTA. Example for a single primer targeting one transcript (MT-CO1). In total, 65 targeted primers are used to enrich mitochondrial transcripts.

**Supplemental Figure 3.**
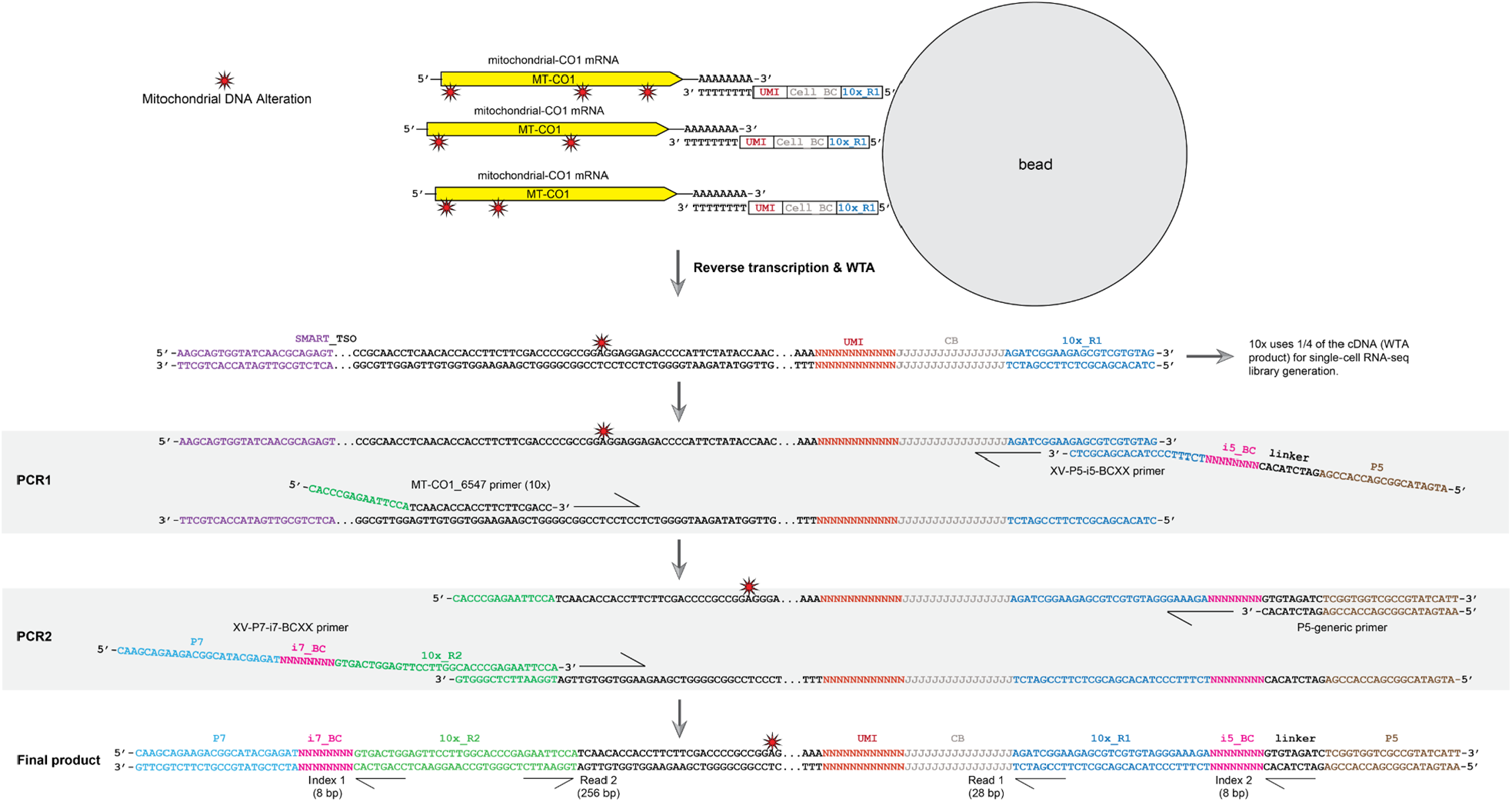
Detailed schematic for MAESTER dial-out PCR protocol from 10x WTA (cDNA). Example for a single primer targeting one transcript (MT-CO1). In total, 65 targeted primers are used to enrich mitochondrial transcripts.

**Supplemental Figure 4.**
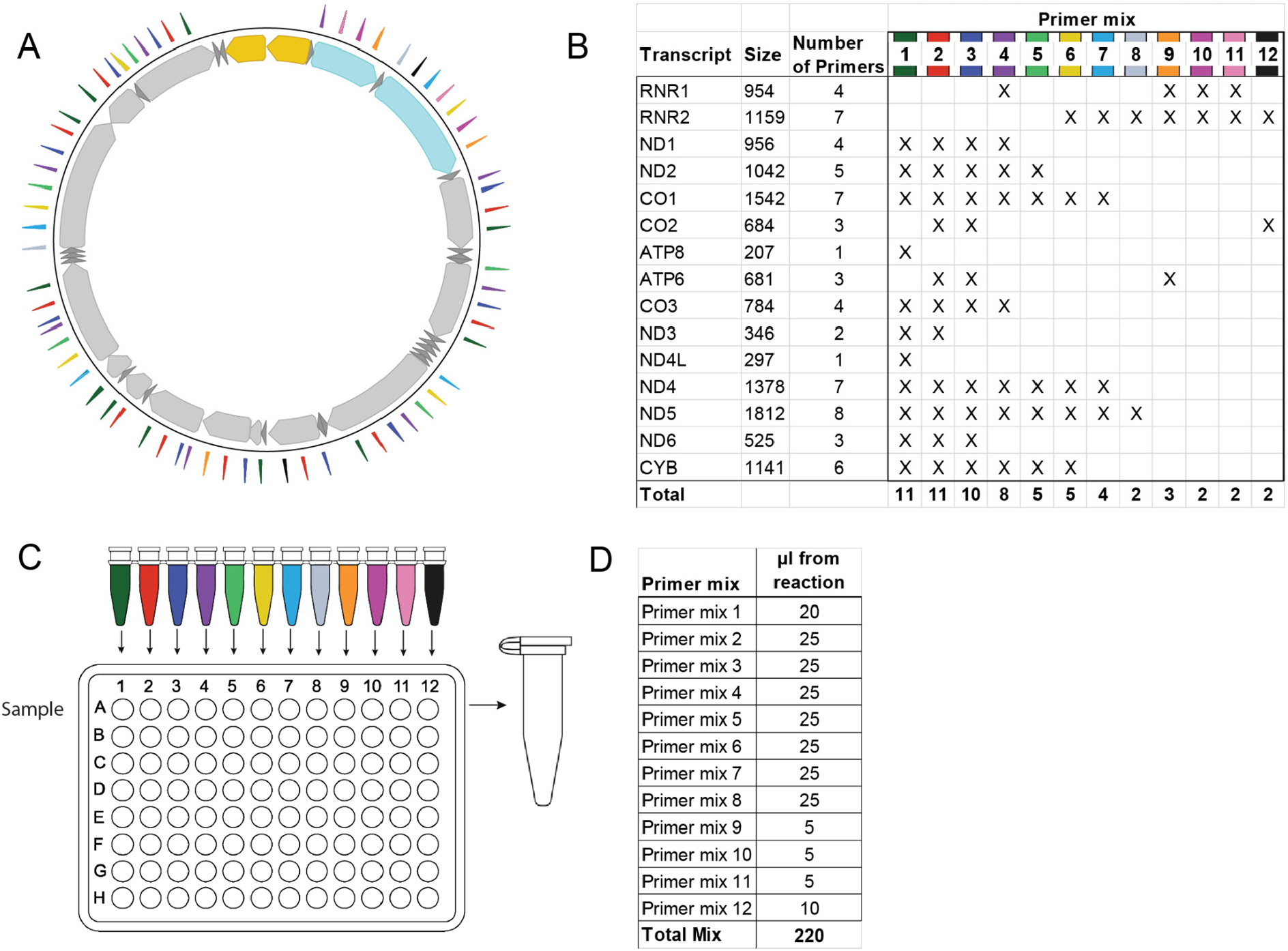
Schematic and protocol for primer mixing. **A**. Diagram depicts the circular mitochondrial genome. The colored triangles indicate where MAESTER primers bind. Colors indicate which primer mix they are in, as detailed in B and C. **B**. Table of 15 transcripts captured by scRNA-seq, with the length of the transcript and number of primers targeting that transcript. Table also indicates the primers for each transcript that are included in each of the 12 primer mixes for PCR1. This experimental design ensures that each transcript is targeted by no more than one primer per mix, that amplicons within the same PCR mix have similar sizes, and that highly expressed amplicons (mainly RNR1 and RNR2) do not overtake the reaction. **C**. Twelve PCR reactions with indicated primer mixes are completed for PCR1. Multiple samples can be processed in a 96 well plate. **D**. After PCR1, the 12 reactions are mixed together according to the volumes in the chart. This mixture is used as a template for PCR2 to generate sequencing libraries.

**Supplemental Figure 5.**
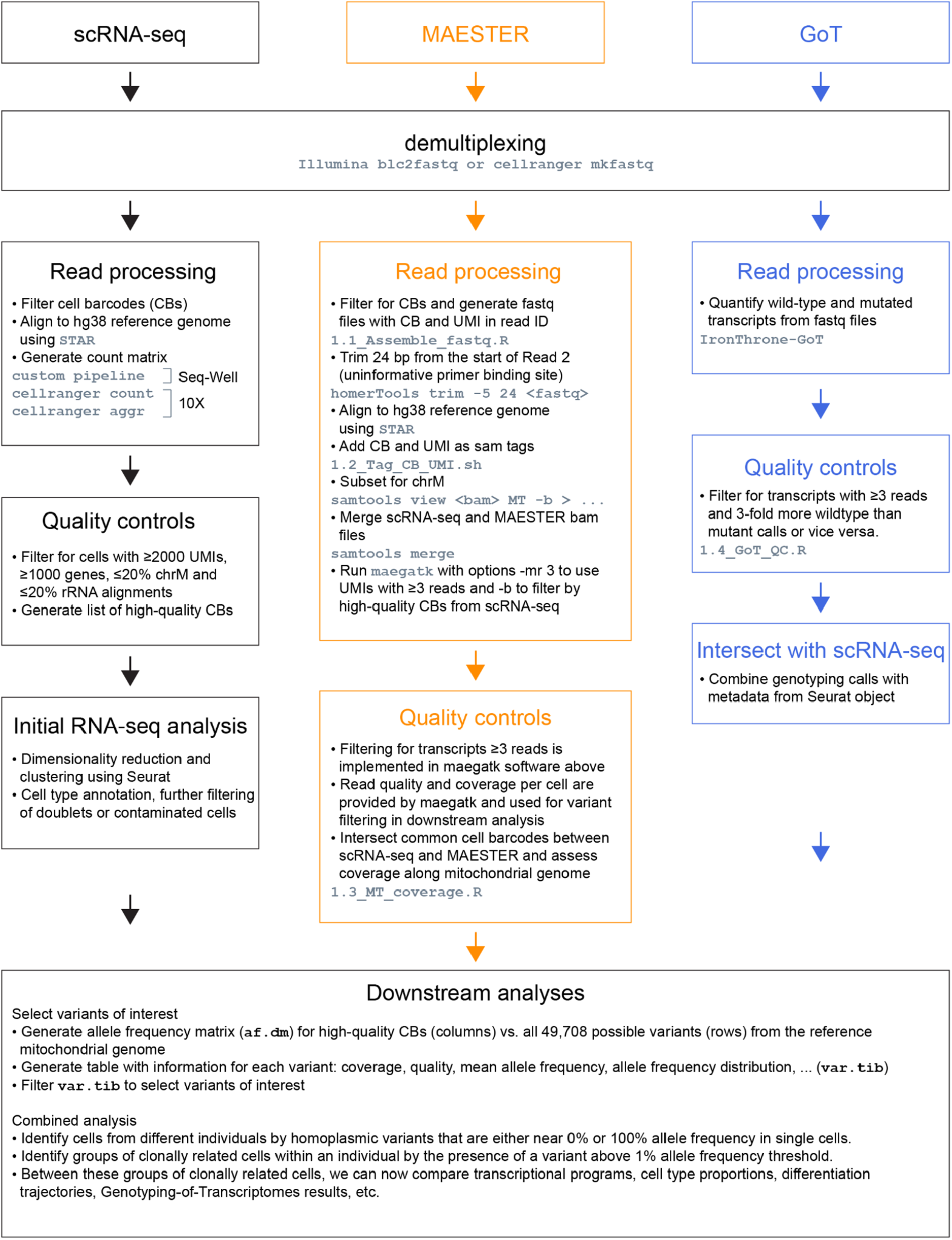
Overview of computational analyses. Flow chart shows the major steps for the integrated analysis of scRNA-seq, MAESTER and Genotyping of Transcriptomes data from the same single cells. Indicated in grey are software tools (bcl2fastq, cellranger, STAR, homerTools, IronThrone-GoT, maegatk) and custom scripts (ending in .sh or .R), all of which are publicly available (see Software and data availability).

**Supplemental Figure 6.**
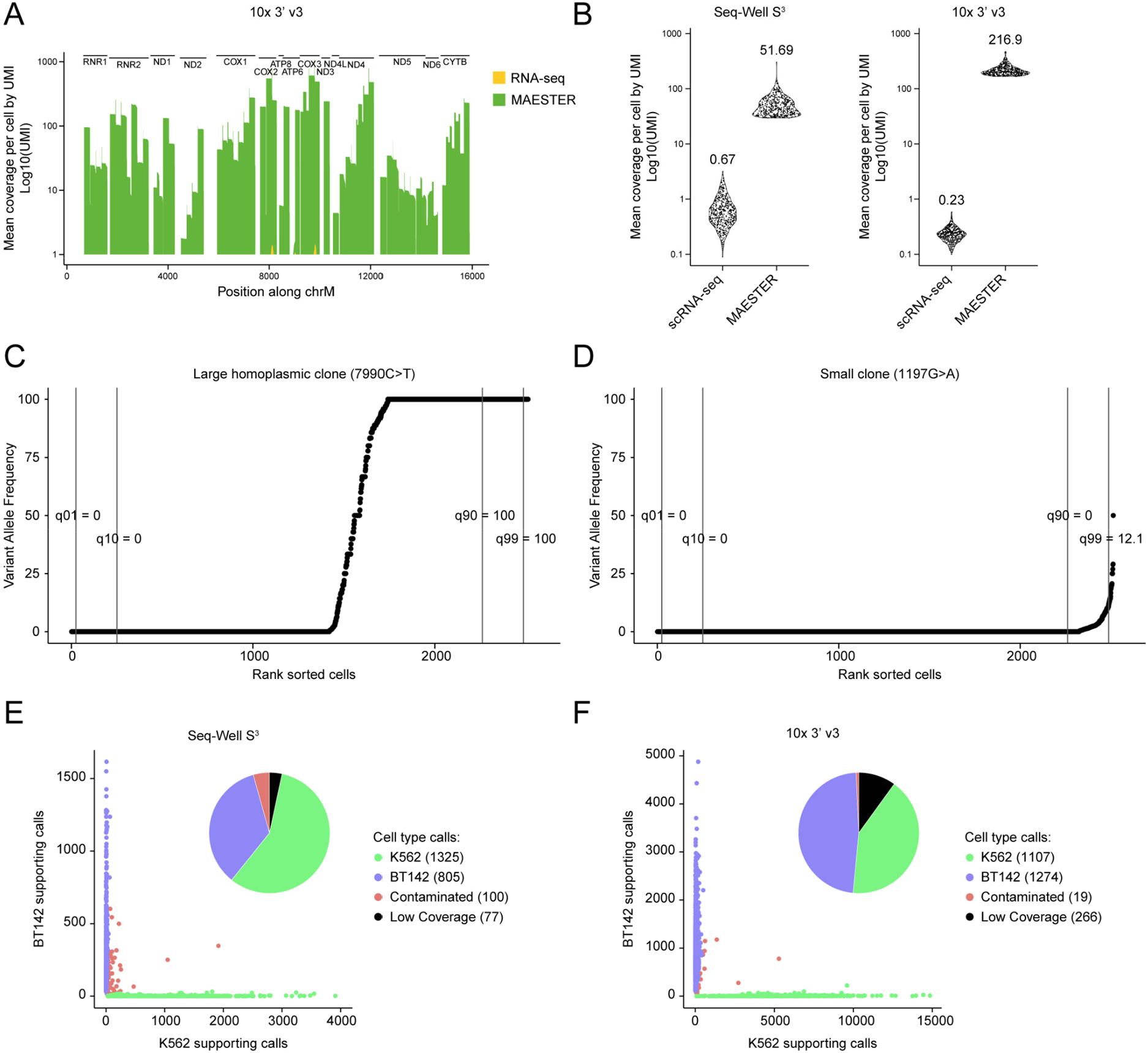
Supporting data for Figure 1. **A**. Bar plot shows coverage of the mitochondrial genome with and without amplification from 10x libraries using MAESTER (see Figure 1C for Seq-Well). Mean coverage of 2,488 K562 and BT142 cells is shown. Each of the transcripts (UMIs) was sequenced ≥3 times. **B**. Sina plots show the mean coverage along all bases of the mitochondrial genome for the top 500 cells (symbols) analyzed by scRNA-seq alone or MAESTER as indicated. The mean for all cells is indicated on top. The difference in MAESTER coverage between Seq-Well and 10x is in part due to different sequencing depths: 200,395,346 and 683,202,846 reads, respectively. **C-D**. Scatter plots depict an important strategy by which informative variants were selected. For each variant the allele frequency across all cells was sorted from low to high, followed by determining the VAF at different quantiles (e.g. the 1st, 10th, 90th and 99th quantile). All 6 (Seq-Well) and 17 (10x) homoplasmic variants that were used to distinguish between cell lines met the following parameters: mean coverage per cell >20, mean read quality >30, 10th VAF quantile of <10% and 90th VAF quantile of >90%. None of the other 49,691 possible variants met these parameters. **E-F**. Scatter plot shows the number of supporting calls (wild-type + mutant) for homoplasmic variants in K562 and BT142 cells. For Seq-Well (E), six variants shown in Figure 1F were used. For 10x (F), seventeen variants shown in Supplemental Figure 7C were used. Of the cells that were classified by mtDNA variants, 2,129/2,130 (100%) were concordant with mRNA clusters for Seq-Well and 2,380/2,381 (100%) were concordant with mRNA clusters for 10x.

**Supplemental Figure 7.**
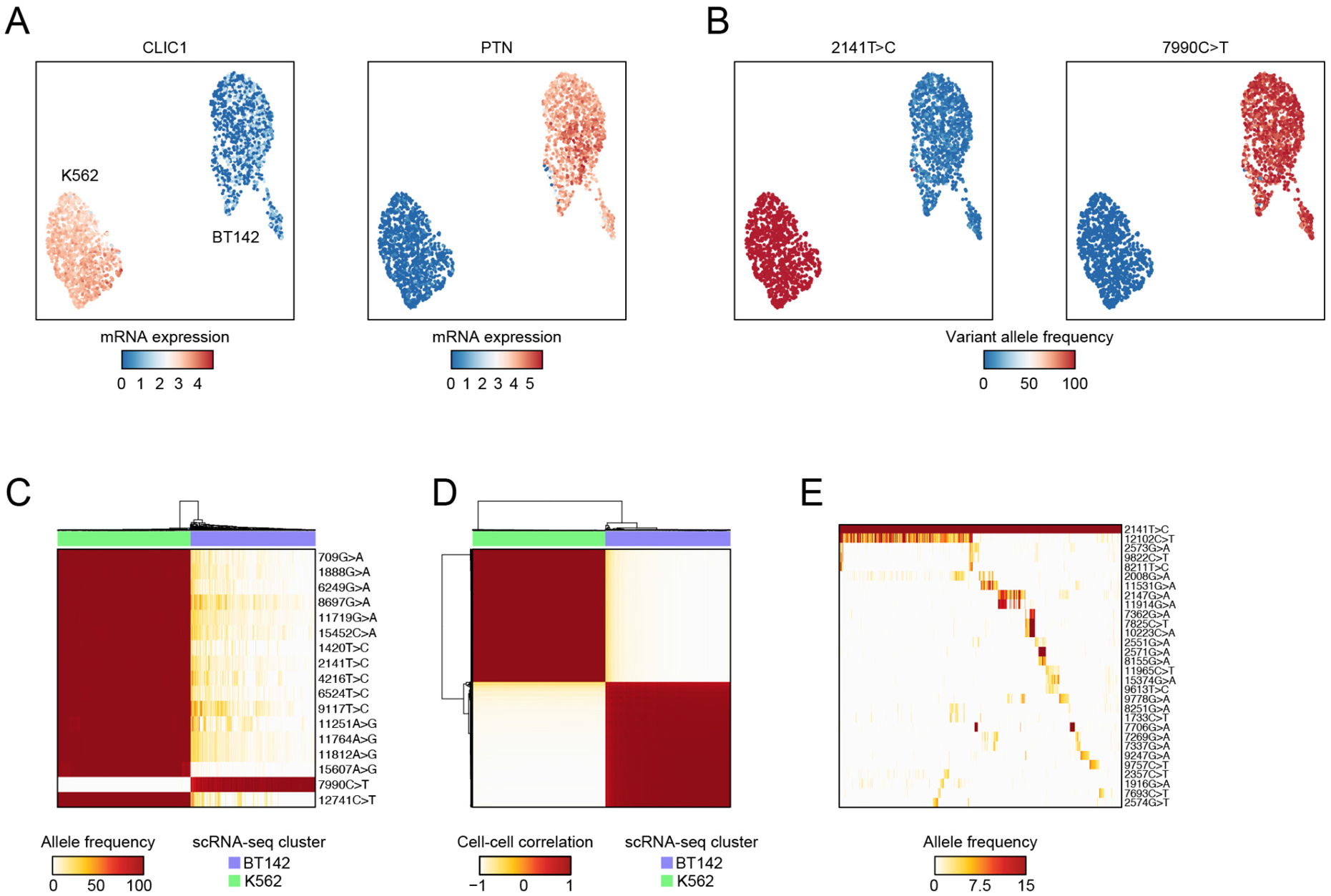
Analysis of 10x cell line mixing experiment. **A-B**. UMAPs show detection of cell-type specific (A) gene expression from scRNA-seq and (B) homoplasmic mtDNA variants from MAESTER. Cluster identity is confirmed by mRNA expression of *CLIC1 (*a K562-specific gene) and *PTN (*a BT142-specific gene). Using MAESTER, we detected the mtDNA variant 2141T>C specifically in K562 cells and 7990C>T specifically in BT142 cells. Only cells with >3-fold coverage of the indicated variants are shown. **C**. Heatmap depicts separation of 2,083 K562 and BT142 cells (columns) based on 17 mtDNA variants detected using MAESTER (rows). Cell type annotation from scRNA-seq is shown on top. **D**. Correlation matrix shows cell similarity based on the allele frequencies of 17 homoplasmic variants (rows and columns depict 2,083 cells). Unsupervised clustering identified two clusters that correlate with cell annotations from scRNA-seq. For C and D, only cells with >3-fold coverage of each homoplasmic variant are shown. **E**. Heatmap shows VAF of 29 mtDNA variants detected by MAESTER (rows) for 811 K562 cells (columns) with informative subclonal variants. Variant 2141T>C is a homoplasmic variant seen in all cells, shown for comparison. Heatmap is organized by clonal structure (Methods). This figure shows analysis of MAESTER with 10x; see Figure 1D-H for similar analysis of MAESTER with Seq-Well.

**Supplemental Figure 8.**
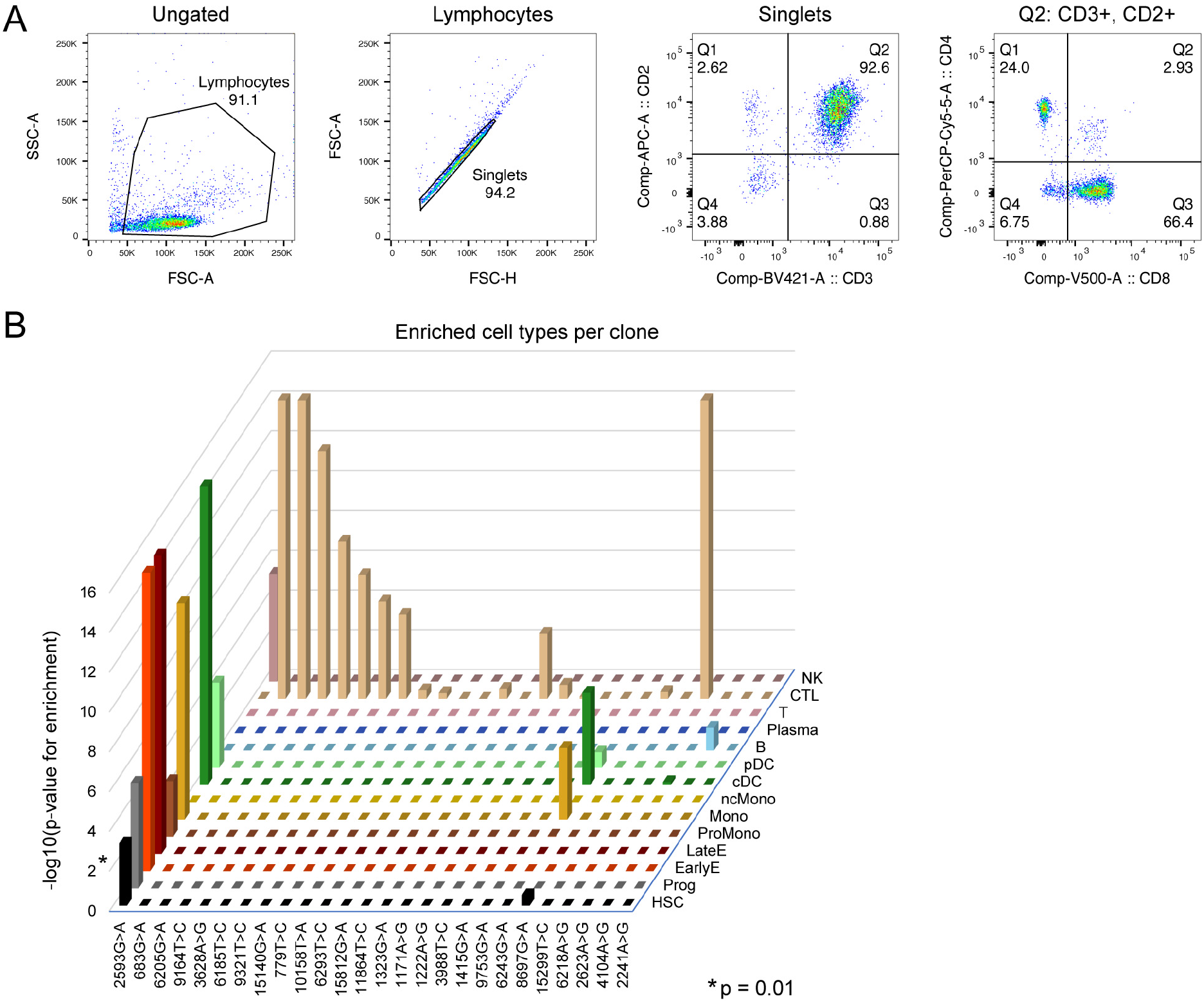
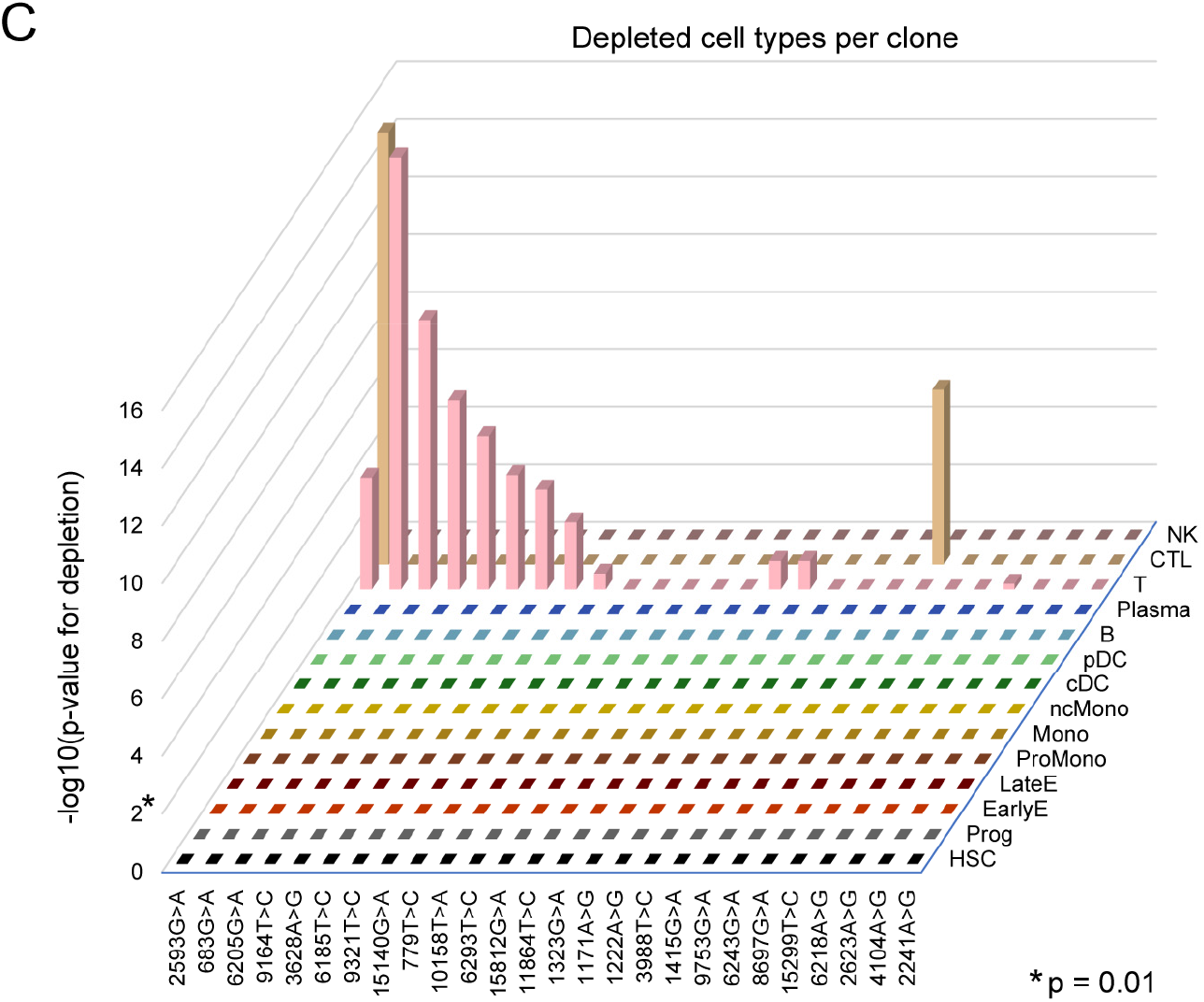
Cell type frequencies in clonal hematopoiesis sample and subclones. **A**. Flow cytometry plots of the same bone marrow aspirate used for 10x sequencing in Figure 2. Single lymphocytes were determined by forward and side scatter. CD2+, CD3+ cells represent T-cells (92.6%). CD2+, CD3-cells represent NK cells (2.6%). Of T-cells, 66% were CD8+. This is consistent with an enrichment of cytotoxic T-cells in the overall sample, in accordance with cell classification by scRNA-seq. **B-C**. Enrichment (B) or depletion (C) of 14 cell types detected in each of the 26 clones compared to expected from the bulk sample, related to Figure 2C. The significance was calculated using the cumulative distribution function of the hypergeometric distribution. *P*-values for enrichment or depletion are shown independently. *P*-values were corrected by Bonferroni correction (n=364).

**Supplemental Figure 9.**
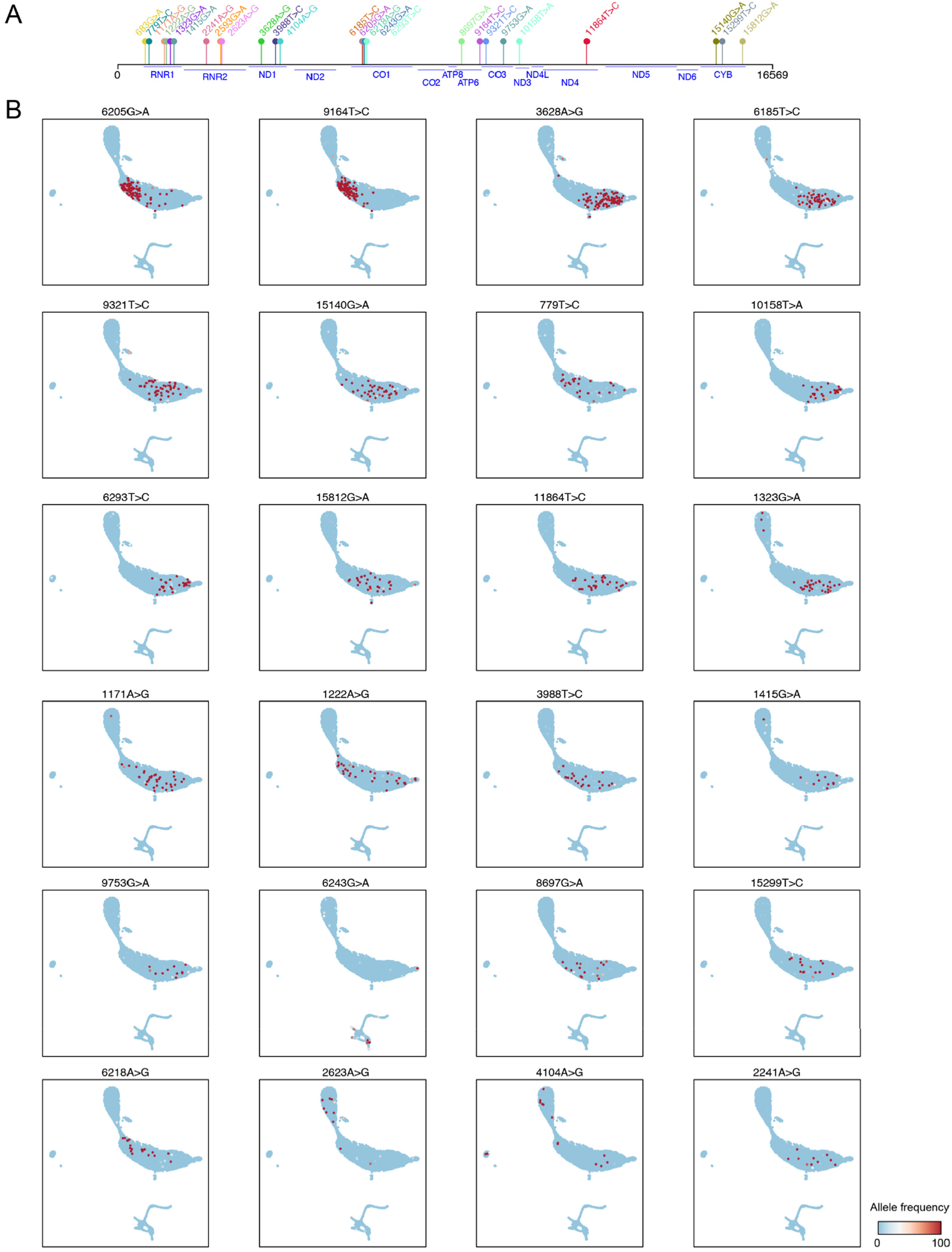
Mitochondrial variants inform clonally related cells in clonal hematopoiesis. **A.** Chart shows the position of subclonal variants that were identified in the clonal hematopoiesis sample, along all 16,569 bases of the mitochondrial genome. At the bottom, the position of genes is indicated in blue. **B.** UMAPs show VAFs per cell for informative variants in Figure 2B. Cell coordinates are the same as in Figure 2A.

## Methods

### Cell lines and culturing

Human chronic myelogenous leukemia K562 cells (ATCC CCL-243) were cultured in RPMI 1640 Medium with GlutaMAX (Gibco 61870127), supplemented with 10% fetal calf serum (FCS) and penicillin-streptomycin. BT142 gliomasphere line^16^ (ATCC ACS-1018) were maintained in culture as previously described^17^. Briefly, neurosphere cultures contain Neurobasal media supplemented with 20 ng/mL recombinant EGF (R and D Systems), 20ng/mL FGF2 (R and D Systems), 1X B27 supplement (Invitrogen), 0.5X N2 supplement (Invitrogen), 3mM L-glutamine, and penicillin/streptomycin. 25% conditioned media was carried over each passage. Cultures were confirmed to be mycoplasma-free via PCR methods.

### Primary human bone marrow

The patient with clonal hematopoiesis in this study consented to all study procedures under a Dana-Farber Cancer Institute IRB-approved research protocol. This patient had a history of leukopenia and thrombocytopenia. The bone marrow sample we analyzed was an aspiration in the context of evaluation for skin-only BPDCN. Histologic evaluation of the concurrent bone marrow core biopsy was normal and did not show involvement by malignant BPDCN cells. Targeted sequencing of the bone marrow aspirate identified multiple alterations indicating clonal hematopoiesis: ASXL1 G642fs in 44.7% of 403 reads, TET2 S792X in 29.5% of 515 reads, TET2 Q1034X in 31.4% of 621 reads, TET2 R1216X in 8.8% of 250 reads and TET2 H1380Y in 6.9% of 567 reads. An aliquot of the bone marrow aspirate was provided for research use. Mononuclear cells were isolated by density centrifugation and cryopreserved with 10% DMSO in liquid nitrogen. Cells were thawed using standard procedures; since viability (independently assessed by Trypan and propidium iodide staining) exceeded 90%, unsorted cells were used for scRNA-seq using the 10x 3’ v3 protocol. A high proportion of cytotoxic T cells was recovered, consistent with expansion of large granular lymphocytes in the peripheral blood of this patient as demonstrated by routine clinical evaluation and confirmatory flow cytometry (Supplemental Figure 8A). This specimen is likely hemodiluted with a contribution from the peripheral blood as the bone marrow core biopsy did not contain this high fraction of T-cells. The scRNA-seq data from this sample is also being utilized in an independent manuscript not involving the MAESTER technique and is under consideration elsewhere (Patient 10, Griffin *et al*., manuscript under review).

### Single-cell RNA-sequencing

For Seq-Well S^3^ experiments, cells were processed as described previously^11^. A complete, updated protocol for Seq-Well S^3^ is hosted on the Shalek Lab website (www.shaleklab.com). Briefly, an array with ∼90,000 nanowells was first loaded with barcoded mRNA capture beads, then 10-15,000 cells were added dropwise onto the surface of the array. After cells were allowed to settle into the wells, the array was sealed with a semi-permeable polycarbonate membrane. Cells were lysed and mRNA transcripts were hybridized to the bead contained within the same well at the polyT sequence of the barcoded oligonucleotides. The beads were then used to generate cDNA via reverse transcription. A second strand synthesis step using a random octamer was performed to recover transcripts in which template switching during reverse transcription was not successful. Whole transcriptome amplification (WTA) PCR was performed and the product underwent a combination of tagmentation and PCR to generate dual indexed sequencing libraries. Libraries were sequenced using a 75 cycle kit on the Illumina NextSeq500 with custom read 1 (CR1P) and custom i5 primers (SW-Ci5P, Supplemental Table 1), 20 cycles for Read 1 (cell barcode or CB + UMI), 56 cycles for Read 2 (transcript sequence), and 2 x 8 bp library barcodes.

For 10x experiments, we used Chromium Single Cell 3′ v3, following all manufacturer’s recommendations. Briefly, 5,000 cells were loaded per well and captured in gel bead-in emulsions (GEMs). Captured mRNAs were reverse transcribed into cDNAs and amplified to generate WTAs. Library construction involves fragmentation, adaptor ligation and a sample index PCR. Libraries were sequenced using a NovaSeq SP 100 cycle kit with 28 cycles for Read 1 (CB + UMI), 91 cycles for Read 2 (transcript sequence) and an 8 bp library barcode.

For the cell line mixing experiments, we used two Seq-Well S^3^ arrays and two 10x 3’ v3 wells, yielding a similar number of cells and data quality. For the primary human sample, we used four 10x 3’ v3 wells.

### Mitochondrial Alteration Enrichment for MAESTER from Seq-well or Drop-seq

We designed an extension of the Seq-Well method for targeted amplification of mtDNA mutations from the WTA product based on a method we initially developed for the detection of somatic mutations^4^. The starting material for this single-cell genotyping method is the product of the Seq-Well WTA reaction (only a fraction of which is used for scRNA-seq). The general method consists of two PCR reactions with a streptavidin bead enrichment in between (Supplemental Figures 1-2). The first PCR reaction serves to add a biotin tag and Nextera adaptor (NEXT) to mitochondrial transcripts while retaining the UMI and CB of the transcripts. The second PCR is used to append Illumina adapters (P5, P7), dual index barcodes to identify the sample, and sequencing primer binding sites.

#### PCR1

We designed biotinylated primers to tile across the entire mitochondrial transcriptome. Primer mixes were made at a 10-fold concentration relative to final concentration in the PCR reaction. Twelve primer mixes were created using 2-11 of these primers at a concentration of 1 µM each. The SMART-AC primer, which is common to all PCR1 reactions, was also included in each primer mix at 10 µM (Figure 1B, Supplemental Figure 4, Supplemental Table 1).

To prepare the template for the MAESTER reaction, WTA products from an individual sample were pooled and diluted to be used at 20 ng in a total volume of 10 µL per reaction. Next, 2.5 µL of the primer mix and 12.5 µL of KAPA HiFi Hotstart ReadyMix (Fisher Scientific KK2602) were added to the template and PCR was performed using the following conditions: initial denaturation at 95C for 3 minutes, followed by 6 cycles of 98C for 20 seconds, 65C for 15 seconds, and 72C for 3 minutes, ending with a final extension at 72C for 5 minutes. There were 12 reactions in total for each sample, as each primer mix is used in a single reaction.

Following amplification, the PCR product is pooled and purified with 0.8x AMPure XP beads to remove primers (Beckman Coulter A63881). Pooling ratios of PCR1 products were empirically determined (Supplemental Figure 4) based on coverage obtained from sequencing to obtain a more equal distribution of reads across the mitochondrial transcriptome. Using Streptavidin-coupled Dynabeads, only biotinylated fragments containing the amplicons of interest are captured (following manufacturer’s instructions, ThermoFisher 60101). Dynabeads/DNA-complex is eluted in 23 µl H2O and used as template for the second PCR.

#### PCR2

To add Illumina adapters (P5, P7), index barcodes to identify the library (i5, i7), and sequencing primer binding sites to the fragments, a second PCR is performed using 23 µl of streptavidin-bound template, with 2 µl of a 5 μM primer mix (N70D_P7_BCXX and N70_P5_BCXX; Supplemental Table 1) and 25 µl PFU Ultra II HS 2xMasterMix (ThermoFisher Q32854). The parameters used for PCR2 are an initial denaturation at 95C for 2 minutes, then 6 cycles of 95C for 20 seconds, 65C for 20 seconds, and 72C for 2 minutes, and then a final extension at 72C for 5 minutes. After the second PCR, the streptavidin beads are magnetized to collect the supernatant, from which DNA is purified with 0.7x AMPure XP beads. After elution in 22 µl TE, the supernatant is transferred to a new tube and saved for sequencing.

The resulting libraries are similar to Seq-Well scRNA-seq libraries but with targeted integration of the NEXT sequencing primer binding site at the region of interest. The libraries were generally 2-10 ng/µl with bands ranging from 250-1000 bp in size. Libraries were sequenced on the Illumina NovaSeq SP 300 cycle kit with CR1P, 20 cycles for Read1, 264 cycles for Read2, and 2 x 8 bp index barcodes.

### Mitochondrial Alteration Enrichment for MAESTER from 10x Genomics

Enrichment for 10x Genomics was very similar to the protocol for Seq-Well or Drop-Seq described above. The main differences are the use of primer sequences specific to 10x and the omission of the biotin enrichment step (Supplemental Figures 1, 3).

#### PCR1

We designed primers to tile across the entire mitochondrial transcriptome. Primer mixes were made at a 10-fold concentration relative to final concentration in the PCR reaction. Twelve primer mixes were created using 2-11 of these primers at a concentration of 1 µM each. A barcoded XV-P5-i5-BCXX primer was included in each primer mix at 10 µM for sample indexing (Supplemental Figure 4, Supplemental Table 2).

To prepare the template for the MAESTER reaction, WTA products from an individual sample were pooled and diluted to be used at 20 ng in a total volume of 16 µL per reaction. Next, 4 µL of the primer mix and 20 µL of KAPA HiFi Hotstart ReadyMix (Fisher Scientific KK2602) were added to the template and PCR was performed using the following conditions: initial denaturation at 95C for 3 minutes, followed by 6 cycles of 98C for 20 seconds, 65C for 15 seconds, and 72C for 3 minutes, ending with a final extension at 72C for 5 minutes. There were 12 reactions in total for each sample, as each primer mix is used in a single reaction.

Following amplification, the PCR product is pooled and purified with 1x AMPure XP beads to remove primers (Beckman Coulter A63881). Pooling ratios of PCR1 products were empirically determined (Supplemental Figure 4D, all volumes multiplied by 1.6) based on coverage obtained from sequencing to obtain a more equal distribution of reads across the mitochondrial transcriptome. After AMPure XP purification, the pooled PCR1 product was eluted in 20 ul H_2_O.

#### PCR2

To add Illumina adapters (P5, P7), index barcodes to identify the library (i5, i7), and sequencing primer binding sites to the fragments, a second PCR is performed using 18 µl of the eluate, with 2 µl of a 5 μM primer mix (P5-generic and XV-P7-i7-BCXX; Supplemental Table 2) and 20 µl KAPA HiFi Hotstart ReadyMix (Fisher Scientific KK2602). The parameters used for PCR2 are an initial denaturation at 95C for 3 minutes, then 6 cycles of 98C for 20 seconds, 60C for 30 seconds, and 72C for 3 minutes, and then a final extension at 72C for 5 minutes. After the second PCR, the DNA is purified with 0.8x AMPure XP beads. The DNA is eluted in 20 µl TE, the supernatant is transferred to a new tube and saved for sequencing.

The resulting libraries are similar to 10x scRNA-seq libraries but with targeted integration at the region of interest. The libraries were generally 2-100 ng/µl with bands ranging from 300-1500 bp in size. Libraries were sequenced on the Illumina NovaSeq SP 300 cycle kit with 28 cycles for Read1, 256 cycles for Read2, and 2 x 8 bp index barcodes. No custom sequencing primers are required.

### Single-cell RNA-seq read processing

For Seq-Well, data was processed as previously described^4^. Briefly, sequencing data was demultiplexed using Illumina’s bcl2fastq2 software (v2.20.0). Read 1 yielded 20 bp reads (12 bp CB and 8 bp UMI), Read 2 yielded 56 bp reads (transcript sequence) and P7 and P5 indices to identify the library were 8 bp each. Reads associated with CBs occurring less than 100 times were removed, and the list of remaining CBs was used to generate Read2 fastq files in which the library barcode, the CB, and the UMI were appended to the read identifier. For 10x, scRNA-seq data was processing using cellranger version 3.1.0 with default settings: cellranger mkfastq to demultiplex into fastq files, and cellranger count to quantify gene expression.

To generate the reference genome, we used hg38 sequences and annotations (v99) from ensembl, with the addition of RNA18S and RNA28S annotations from UCSC. Annotations were filtered using cellranger (version 3.1.0) mkgtf with recommended attributes as well as the gene biotypes gene_biotype:Mt_rRNA and gene_biotype:rRNA. The reference genome was then generated with cellranger mkref which includes STAR indexing. To align Seq-Well scRNA-seq data to this reference, we used STAR version 2.6.0c with the options --outSAMtype BAM SortedByCoordinate and --quantMode TranscriptomeSAM. To align 10x data scRNA-seq data to this reference, we used cellranger count which implements STAR.

### MAESTER read processing

MAESTER fastqs include Read 1 encompassing the CB and UMI (20 bp for Seq-Well, 28 bp for 10x), Read 2 covering mitochondrial transcript sequences (264 bp for Seq-Well, 256 bp for 10x), and 2 x 8 bp for dual-indexed library barcodes. We used Illumina bcl2fastq for demultiplexing with both indices. Reads associated with CBs occurring less than 100 times were removed, and the list of remaining CBs was used to generate Read2 fastq files in which the library barcode, the CB, and the UMI were appended to the read identifier. We trimmed the first 24 bp from these fastqs using homerTools to avoid using the primer binding sequence for variant calling. Next, we aligned the fastq files with STAR (--outSAMtype BAM SortedByCoordinate) to the same hg38 reference genome we used for scRNA-seq alignment above. More than 90% of MAESTER reads aligned to chrM. See Supplemental Figure 5 for an overview of these procedures.

### maegatk - mitochondrial genome variant calling

To facilitate the analysis of MAESTER data, we developed maegatk, a Python package, as an extension of our previously described mgatk pipeline^8^. Importantly, maegatk specifically handles technical biases implicit in high-throughput scRNA-seq to facilitate the robust identification of mtDNA variants. First, maegatk takes inputs of a single-cell .bam file following the 10x Genomics SAM tag conventions, a valid list of CBs, and more than 20 customizable command-line arguments. Next, the software collapses duplicate reads based on UMI, start position, and CB. Unlike most existing variant calling pipelines, including GATK and mgatk that select a representative read (based on highest mean base quality score), maegatk identifies the most likely consensus nucleotide across sequencing read replicates via the CallMolecularConsensusReads (v1.1) tool from fgbio. Further, maegatk provides a --min-reads command line argument that specifies the minimum number of sequence reads needed for a UMI to be considered for variant calling. This workflow minimizes artifacts due to PCR amplification and sequencing error compared to the standard Picard MarkDuplicates. After consensus read deduplication, per-cell, per-position nucleotide counts are enumerated and used for downstream analysis.

For use in our analysis with maegatk, we ensured that all bam files contained the CB and UMI SAM tags according to 10x conventions (CB:Z and UB:Z). We selected reads aligning to chrM and merged bam files from scRNA-seq and MAESTER (Aligned.sortedByCoord.out.bam from STAR or possorted_genome_bam.bam from cellranger). We generated a list of CBs by intersecting CBs with ≥100 alignments to chrM and high-quality CBs from scRNA-seq. Maegatk was then executed with the options --input merged.bam --mito-genome chrM.fa --barcodes CBs.txt --min-reads 3. The last option specifies only UMIs with three reads are used to increase confidence in variant calls. Upon completion, maegatk saves mutation calls as a maegatk.rds file in the SummarizedExperiment format^18^ for convenient intersection with other modalities and downstream analysis in R.

### Single-cell RNA-seq clustering and cell type annotation

For Seq-Well and 10x scRNA-seq data alike, we filtered for cells with ≥2,000 UMIs, ≥1,000 genes, ≤20% alignment to rRNA genes, and ≤20% alignments to genes on chrM. Genes from chrX and chrY were removed from the count matrix. Next, we used Seurat version 3.2.2 for standard scRNA-seq processing steps including the functions NormalizeData, FindVariableFeatures, ScaleData, and RunPCA (similar to https://satijalab.org/seurat/archive/v3.2/pbmc3k_tutorial.html)^19^. We implemented graph based clustering with FindNeighbors with 6 PCA dimensions and FindClusters with a resolution of 0.05-0.1, as we only aimed to distinguish K562 and BT142 cells. We determined to top 10 cell type specific genes by fold change using the FindAllMarkers function with only.pos = TRUE, min.pct = 0.25 and logfc.threshold = 0.25. For Seq-Well, this yielded *FTL, PRAME, NPM1, LDHA, ATP5MC3, PABPC1, ENO1, HBG2, RANBP1* and *HSP90AB1* for K562 and *MET, PTN, PTPRZ1, BCAN, MAP2, TUBA1A, MAP1B, DBI, SOX2* and *FAM3C* for BT142. For 10x, this yielded *CLIC1, ATP5MC3, FTL, PRDX1, MYL12A, LDHA, ENO1, UQCRH, PRAME* and *RANBP1* for K562 and *PTN, BCAN, SOX2, TUBA1A, CLU, PTPRZ1, DBI, NOVA1, TSC22D1* and *MARCKS* for BT142.

For cell line mixing experiments, we used decontX from the R package celda version 1.5.6 to remove cells with high ambient RNA^20^. We supplied the decontX function with the count matrix to calculate per-cell contamination scores. We also scored each cell for both cell type specific gene signatures using the Seurat function AddModuleScore. Finally, we excluded cells that exceeded a contamination score of 0.05 *and* had a high module score for both cell type specific signatures. For Seq-Well, this removed 218/2525 (8.6%) of cells, with 1387 K562 and 920 BT142 cells remaining. For 10x, this removed 112/2778 (4.0%) of cells, with 1310 K562 and 1356 BT142 cells remaining.

For the clonal hematopoiesis sample, we used the cell type annotations that were previously established using a Random Forest Classifier based on healthy donor populations (Griffin *et al*., manuscript under review). Filtering for high-quality cells, dimensionality reduction and removal of cells with ambient RNA were performed using similar procedures as the cell line mixing experiments except decontX. All scripts are available on Github.

### Identification of informative mtDNA variants

We took similar approaches to select informative mtDNA variants to (1) distinguish K562 and BT142 cells by homoplasmic variants, (2) reveal clonal structure within the K562 population and (3) determine clonally related cells in the clonal hematopoiesis sample. From the output of the maegatk software, we calculated an allele frequency matrix of all possible variants (rows) and cells (columns). The total number of possible variants is 3×16,569+1=49,708 because the mitochondrial genome (NC_012920) is 16,569 bp with three possible variants each, except base 3,107, which has four possible variants (A, C, T, G) because the reference is N. Next, we generated a table with features for every variant: the mean allele frequency, mean coverage, mean quality score, and the VAF in quantiles of rank sorted cells (Figure S6C). This allowed us to select informative variants by applying filters such as mean coverage >20, mean quality >30, VAF of <10% in the bottom 10% of cells and VAF of >90% in the top 10% of cells. Variants selected as such were highly enriched for transitions vs. transversions, as expected (>93%). Additional filters to remove artifacts were included for subclone identification, for example, variants informing K562 subclones should be absent in BT142 cells. The classification of K562 and BT142 cells by mtDNA variants was determined by the sum of calls at all homoplasmic variants for either cell line (Supplemental Figure 6E-F). In addition to mutant calls, wild-type calls were also used for cell line classification.

Having identified informative variants, we assessed their VAFs in single cells by UMAP visualization (Figures 1E, 2D, Supplemental Figures 7B, 9). For all subclonal analyses, we used a VAF threshold of 1% to consider a cell positive for a mtDNA variant. We combined highly correlated variants together into groups or clones (e.g. 10158T>A and 6293T>C, Figure 2B). To visualize subclonal structures in VAF heatmaps (Figure 1H, Figure 2B, Supplemental Figure 7E), we first sorted clones by the number of variants and the number of cells each contained, and then sorted the cells (columns) within each clone as follows (starting with the smallest clone): for clones identified by a single variant, we sorted the cells from high to low VAF; for clones identified by multiple variants, we either sorted by the sum of the VAFs, or clustered the cells by Pearson correlation. For downstream analyses, we classified each cell by the presence of informative variants and intersected this information with gene expression, GoT results and other metadata such as cell type classification and pseudotime ordering.

### Cytotoxic T lymphocyte correlation

To investigate clonally related CTLs, we assessed correlation of gene expression. First, we used the Seurat function FindVariableFeatures to identify the 100 most variable genes between all 6,130 CTLs in the sample: *S100B, PTGDS, IL7R, LTB, TRDC, GNLY, GZMK, STMN1, KLRB1, PCLAF, H1-3, H1-4, KLRC1, TRBV28, TYMS, TK1, CLSPN, LGALS1, CCDC168, TRDV2, CCL4L2, TYROBP, IFI27, TRAV17, MKI67, FAM111B, CCR7, TRBC1, ACTB, CTSG, NEFL, TRGV9, CENPF, H4C3, RRM2, FCER1G, TOP2A, E2F1, TRGC1, COTL1, CEP55, MAL, H1-2, TRGC2, TRBV6-2, IFI44L, SCGB3A1, CCL4, SPON2, SYNE2, IGKC, SELL, DLGAP5, SLC4A10, TUBA1B, TSHZ2, TMEM155, H1-10, FGFBP2, ACTG1, GZMB, TRBV18, RTKN2, CEBPD, KLRC2, CENPU, JUN, ZNF683, TRAV5, HMMR, RCAN3, RSAD2, DST, TRBV23-1, ISG15, PRSS57, TTN, HLA-DRA, LEF1, LCP1, VIM, IFI6, ME1, AKR1C3, IGFBP2, MCOLN3, KRT1, DPP4, STAT1, ASF1B, F5, UBE2C, MX1, LMNA, HMGB2, TRGV10, BCL11A, P2RY14, SLC22A17* and *THEMIS*. Then, we subsetted the normalized gene expression matrix for these 100 variable genes and for all 1,044 CTLs from the 23 subclones. The Pearson correlation between cells of this subsetted gene expression matrix is shown in Figure 2E. Next, we calculated the Pearson correlation of every CTL in a particular clone with every other CTL in that clone, and took the average (intraclonal correlation). We also calculated the Pearson correlation of every CTL in a particular clone with every CTL *not* from the same clone, and took the average (interclonal correlation). These values are shown in Figure 2F.

### Genotyping of Transcriptomes analysis

To analyze reads from the GoT analysis for the clonal hematopoiesis sample, we utilized IronThrone-GoT Version 1.0 (https://github.com/landau-lab/IronThrone-GoT)^3^. We provided the IronThrone-GoT function with fastq files and a whitelist of CBs for 10x 3’ v3 scRNA-seq. From the summTable, we selected high-confidence transcript calls by filtering for UMIs that were sequenced ≥3x with ≥3x more wild-type than mutant calls or vice versa. We then generated a table with CBs, the number of wild-type transcripts and the number of mutant transcripts per cell. These calls were intersected by CBs with cells of different subclones, as shown in Figure 2H.

### Pseudotime analysis

We analyzed the myeloid differentiation trajectory and assigned pseudotime values to cells using the R slingshot library^14^. We selected all HSCs, progenitors, promonocytes, monocytes and non-classical monocytes from the clonal hematopoiesis sample and provided the slingshot function with the UMAP coordinates. The predicted trajectory (black curve) and assigned pseudotime values are shown in Figure 2I, which is an enlargement of Figure 2A that also includes cDC, pDC and erythroid cells (in grey). The same pseudotime values are used for the horizontal axis of Figure 2J.

### Code or programs used to generate figures

Geneious Prime with Primer3 was used for primer design. Read processing was performed using command line tools including Samtools, cellranger, homerTools, STAR and IronThrone-GoT. Quality controls and downstream analyses were performed with R version 3.6.1 with RStudio version 1.2.5042 and the tidyverse version 1.3.0 collection of packages including ggplot2 version 3.3.2. We used SummarizedExperiment for maegatk output, celda for decontX and slingshot for trajectory analyses as described in the provided Github scripts. We also used GraphPad Prism, Microsoft Excel and Adobe Illustrator for additional statistical analyses and visualization.

## Software and data availability

Read files (fastq) will be deposited in GEO (https://www.ncbi.nlm.nih.gov/geo/). Single-cell gene expression matrices, mtDNA variant calls and GoT results are available at https://vangalenlab.bwh.harvard.edu/resources/maester-2021/ (please contact the corresponding author for the password). maegatk version 0.1 is available at https://github.com/caleblareau/maegatk. The computational analysis is described at https://github.com/vangalenlab/MAESTER-2021.

